# A chemically defined biomimetic surface for enhanced isolation efficiency of high-quality human mesenchymal stromal cells under xeno-/serum-free conditions

**DOI:** 10.1101/2021.12.10.472047

**Authors:** Kristina Thamm, Kristin Möbus, Russell Towers, Stefan Baertschi, Richard Wetzel, Manja Wobus, Sandra Segeletz

## Abstract

Mesenchymal stromal cells (MSCs) are one of the most frequently used cell types in regenerative medicine and cell therapy. Generating sufficient cell numbers for MSC-based therapies is constrained by: 1) their low abundance in tissues of origin, which imposes the need for significant ex vivo cell amplification, 2) donor-specific characteristics including MSC frequency/quality that decline with disease state and increasing age, 3) cellular senescence, which is promoted by extensive cell expansion and results in decreased therapeutic functionality. The final yield of a manufacturing process is therefore primarily determined by the applied isolation procedure and its efficiency in isolating therapeutically active cells from donor tissue. To date, MSCs are predominantly isolated using media supplemented with either serum or its derivatives, which pose safety and consistency issues. To overcome those limitations while enabling robust MSC production with constant high yield and quality, we developed a chemically defined biomimetic surface coating, called isoMATRIX, that facilitates the isolation of significantly higher numbers of MSCs in xeno-/serum-free and chemically defined conditions. The isolated cells display a smaller cell size and higher proliferation rate than those derived from a serum-containing isolation procedure and a strong immunomodulatory capacity. In sum, the isoMATRIX promotes enhanced xeno-, serum-free, or chemically defined isolation of human MSCs and supports consistent and reliable cell performance for improved stem cell-based therapies.

## Introduction

Adult stem cells, also called somatic stem cells, offer great promise for the treatment of intractable diseases and disorders. They are isolated from perinatal or adult tissues and bear the advantage of restricted lineage potential that reduces the risk of tumorigenesis compared to pluripotent stem cells^1,2^. An attractive source of somatic stem cells is mesenchymal stromal cells (MSCs), a population of multipotent progenitor cells capable of differentiating into osteogenic, adipogenic, and chondrogenic lineages. Their diverse therapeutic potential, safety, and ability to self-renew with high proliferative capacity make them especially valuable for regenerative medicine and cell therapy^3^. MSCs can ameliorate diseases, support engraftment, and modulate the host immune response mainly via the secretion of paracrine factors, such as cytokines and growth factors, or extracellular vesicles^4,5^. The success of such innovative cell therapies depends on the availability of stem cell sources and the efficacy of isolation and expansion techniques to yield sufficient amounts of therapeutically active cells.

MSCs can be isolated from various tissues such as bone marrow, adipose tissue, umbilical cord, or deciduous teeth. The number of stem cells varies considerably depending on the tissue of origin. Bone marrow for example harbors the lowest amount of MSCs (1 to 30 cells/mL to 317,400 cells/mL), followed by adipose tissue (4,737 cells/mL of tissue to 1,550,000 cells/mL) and umbilical cord tissue (10,000 cells/mL to 4,700,000 cells/cm of umbilical cord)^6^. Comparative studies show that MSCs from bone marrow, adipose tissue, and umbilical cord share a common phenotype, differentiation potential, and the expression of many cell surface markers. However, they noticeably differ in gene expression patterns, proliferation rate, and therapeutic efficacy^7–9^.

The conventional method to isolate MSCs involves their selection via non-specific adhesion to plastic cell culture ware in serum-containing medium over several passages, which introduces early cellular heterogeneity and might reduce isolation efficiency. In addition, the transfer of MSCs from their native microenvironment to the nutrient-rich artificial culture conditions induces a metabolic shift from glycolysis towards oxidative phosphorylation resulting in high metabolic heterogeneity and accelerated senescence^10^. The use of fetal bovine serum (FBS) introduces undesirable variability due to its poorly-defined composition, leading to inconsistent lot-to-lot performance and appreciable risk of contamination and adverse immunological reactions against xenogeneic components. To eliminate animal-derived components during isolation and ex vivo expansion of human MSCs for clinical applications, human serum, and platelet lysate are frequently used as growth supplements and reported to support enhanced proliferation of MSCs^11–13^. However, their quality is affected by a several factors such as preparation procedure, donor age or blood profile, and heparin levels (in platelet lysate to prevent clotting)^14,15^. Furthermore, concerns regarding the availability of large quantities required for clinical applications and immunological responses, and transmission of human infections have been raised.

Growth supplements, isolation procedures, and culture conditions can affect the phenotype, stemness, and differentiation potential of MSCs, thereby limiting their clinical applicability^16–18^. Thus, identifying optimal conditions for isolation and in vitro expansion of MSCs is crucial for the development of advanced cell therapies. Optimization includes the adjusting of medium composition, culture substrate, cell seeding density, O_2_ and CO_2_ concentration, pH, and temperature for more defined and reproducible results. Xeno-/serum-free media are preferably used for MSC isolation and in vitro expansion due to their safety, consistency, and capacity to support MSC proliferation. These media often lack adhesion molecules required for cell attachment and therefore need to be complemented by extracellular matrix proteins such as collagens, fibronectin, vitronectin, or laminin. Since many of these biomaterials are of animal origin, they share the aforementioned disadvantages of batch-to-batch variabilities and concerns regarding pathogen contamination and immune responses after cell implantation.

We recently developed a chemically defined biomatrix for the in vitro culture of MSCs, named myMATRIX MSC, that enables a more reliable and reproducible expansion. This matrix is formed by sulfated glycosaminoglycan (GAG) interactions with a four-arm polyethylene glycol-modified with a biofunctional peptide^19^. It promotes cell adhesion and expansion using different media compositions and supports the long-term culture of MSCs in a serum-free medium with robust growth and high viability while maintaining their differentiation and immunomodulatory capacity, characteristic cell morphology, and the expression of key stemness markers^20^.

Analogous to our previous strategy, we have now developed a biomatrix specifically designed for the isolation of MSCs in xeno-/serum-free conditions, called the isoMATRIX. It is composed of dextran sulfate and an extracellular matrix (ECM) protein-derived peptide-conjugate and increases the isolation efficiency of MSCs from bone marrow aspirates and subcutaneous fat. It also reduces biological variations due to its chemically defined nature and compatibility with xeno-/serum-free as well as chemically defined medium. The isolated MSCs show characteristic phenotype, high viability, high proliferation, differentiation potential, and strong immunomodulatory capacities.

## Results

### Xeno-/serum-free isolation of BM-MSCs with isoMATRIX results in a 4-fold increase in isolated cell number and significant decrease in cell size

We previously developed myMATRIX MSC, a surface coating optimized for MSC expansion, which showed suboptimal support during MSC isolation from bone marrow aspirates (data not shown). To identify a new biomatrix that can aid MSC isolation, we selected 12 candidates, previously identified from our in-house screening tool (screenMATRIX), that supported efficient MSC expansion^20^. In preliminary isolation tests, the use of a biomatrix composed of dextran sulfate and a fibronectin-derived peptide-conjugate resulted in the best cell morphology and highest isolated cell number (data not shown). Hence, this biomatrix was chosen for further development and tested for its ability to support the isolation of MSCs using xeno-/serum-free and chemically defined medium. To isolate human bone marrow-derived MSCs (BM-MSCs), mononucleated cells were seeded in duplicates into uncoated 6-well plates using serum-containing medium (TCP control) or in well plates coated with the new biomatrix, called isoMATRIX, in combination with a xeno-/serum-free medium. After 24 h of incubation, non-adherent cells were washed off, and fresh medium was added. On day 4 post-isolation, the number of attached cells was manually counted in 10×13 tile images. An increase of approximately 30% in isolated cell numbers was detected when using the isoMATRIX compared to the control condition (Fig. 1A). This increase led to a significant 4-fold increment in the number of isolated MSCs on the isoMATRIX compared to TCP control 11-14 days post-isolation (Fig. 1B, average ± standard deviation: 131.737 ± 184.801 on TCP; 543.889 ± 474.790 on isoMATRIX). In addition, cells showed high viability (Fig. 1C) and the typical spindle-shaped, fibroblast-like morphology (Fig. 1D) after isolation with the isoMATRIX. Importantly, we were not able to isolate any cells using uncoated or fibronectin-coated culture vessels.

**Figure 1.**
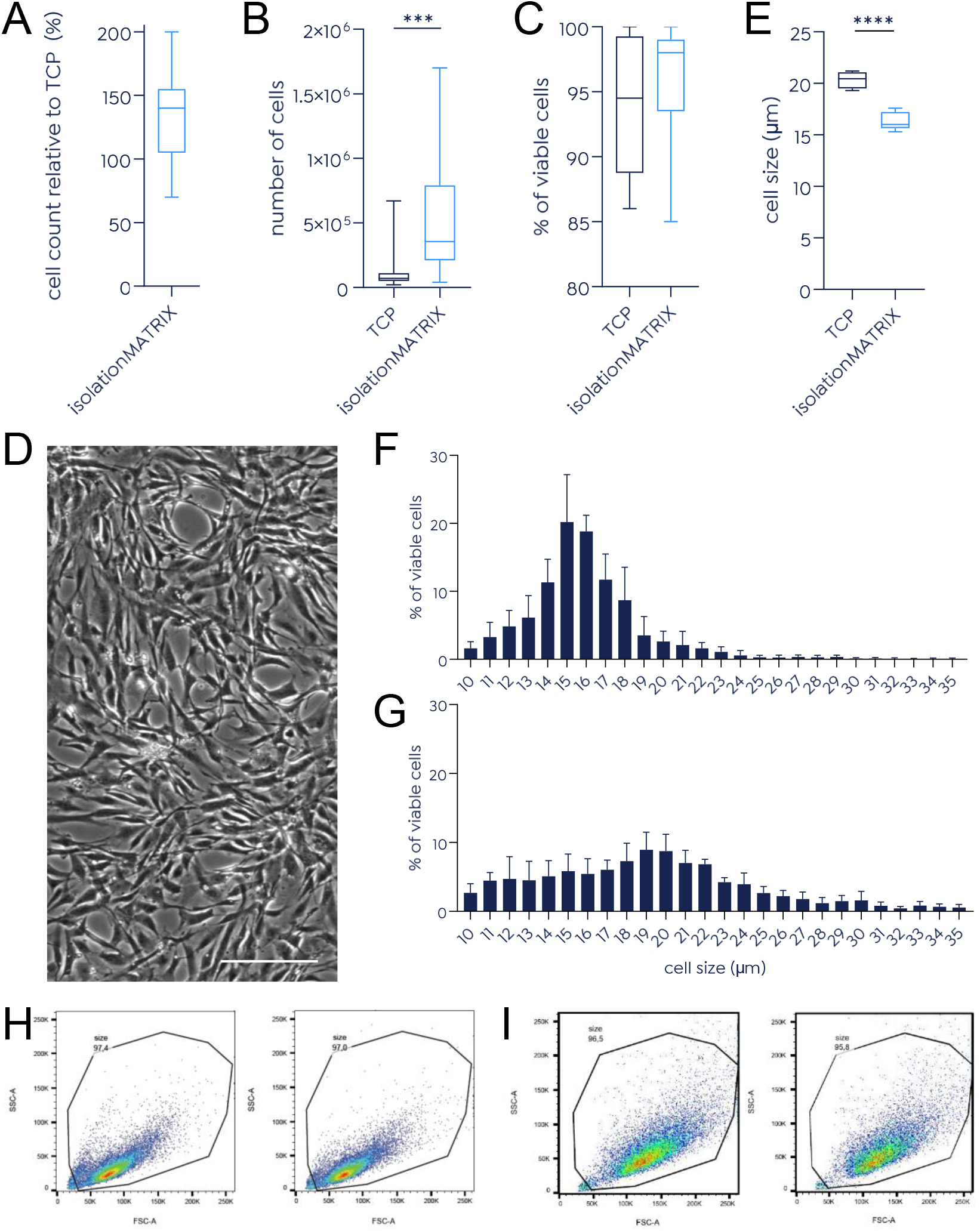
Increased isolated cell numbers, high viability and decreased cell size of BM-MSCs isolated using the isoMATRIX in xeno-/serum-free conditions. Mononucleated cells were seeded in duplicates into uncoated (TCP) or isoMATRIX-coated 6 well plates using serum-containing medium and xeno-/serum-free medium, respectively. After 24h, non-adherent cells were washed off and fresh medium was added. On day 4 after isolation, the cell number was determined by manual counting of 10×13 tile images using Fiji software (A). After 11-14 days, cells were harvested and the number (B), viability (C), morphology (D) and size (E) of isolated MSCs were determined. Cell size distribution of BM-MSCs isolated in xeno-/serum-free medium in combination with the isolationMATIRX (by cell counter (F) or by flow cytometry (H)) or in serum-containing medium without any coating (by cell counter (G) or by flow cytometry (I)) was analyzed. N = 10. Scale bar, 200 μm. *** = *p* < 0.005, **** = *p* < 0.0001

We and others have previously reported that BM-MSCs expanded in serum-free medium show a smaller cell size than MSCs expanded in serum-containing medium^20,21^. Similarly, cells isolated using the isoMATRIX in combination with xeno-/serum-free medium were significantly smaller (16 ± 0.8 μm) compared to those isolated under serum-containing conditions (20 ± 0.8 μm; Fig. 1E). The high growth rates of isoMATRIX-derived MSCs support the concept that small cell size indicates high proliferative activity and low senescence ^22,23^. Moreover, the isolation of cells using the isoMATRIX with xeno-/serum-free medium resulted in a more homogeneous cell population with a cell size of 10-22 μm for most of the cells (96.4% of total counted cells; Fig. 1F) compared to control cells of 10-30 μm (96.7% of total counted cells; Fig. 1G). This increase in homogeneity of cells isolated using the isoMATRIX is also reflected in the flow-cytometry-based size distribution shown in Fig. 1 H.

### BM-MSCs isolated under xeno-/serum-free conditions maintain characteristic expression profiles of cell surface antigens

The expression of specific cell surface antigens is an essential criterion for the characterization of human BM-MSCs. According to the International Society for Cellular Therapy (ISCT), BM-MSCs isolated and expanded in serum-containing media on a plastic surface should express CD73, CD105, and CD90 in at least 95% of the cell population. In contrast, they must not express the hematopoietic markers CD45, CD34, CD11b or CD14, CD79α or CD19 and HLA-DR (<2% positive)^24^. In addition, other cell surface markers such as CD44, CD146, Stro-1, or CD271 have been used to enrich a human MSC population with trilineage differentiation and colony-forming abilities^25^.

We therefore examined the expression of ten cell surface makers (CD73, CD105, CD90, CD44, CD146, CD166, CD11b, CD14, CD45 and CD34) by flow cytometry using cells of passage 2 and 3 (Fig. 2). Almost all cells (99 ± 1.6%) isolated in xeno-/serum-free medium using the isoMATRIX were positive for CD73, CD105, CD90 and CD44. In addition, they showed high levels of CD146 (92 ± 6%) and CD166 (78 ± 11%) (Fig. 2A) and did not express CD34, CD11b, CD14 and CD45 (1.0 ± 0.8%, 0.8 ± 0.2%, 0.6 ± 0.3% and 0.2 ± 0.3%, respectively; Fig. 2B). As postulated by the ISCT, >95% of TCP-isolated cells expressed CD73, CD105 and CD90 (Fig. 2C). 98.9% also expressed CD44. These cells showed a similar expression level of CD146 (93.5 ± 6.4%) but a higher expression of CD166 (98.7 ± 0.9%) compared to cells isolated under xeno-/serum-free conditions. In TCP-isolated cells, the expression of hematopoietic markers CD34, CD11b, CD14 and CD45 was higher compared to isoMATRIX-derived BM-MSCs (3.1 ± 1.3%, 2.5 ± 1.3%, 3.1 ± 3.8% and 1.4 ± 1.0%, respectively), often exceeding the 2% limit postulated by the ISCT (Fig. 2D). Thus, BM-MSCs isolated under xeno-/serum-free conditions with the isoMATRIX fulfill the requirements of international guidelines with a better delineation from hematopoietic stem cells.

**Figure 2.**
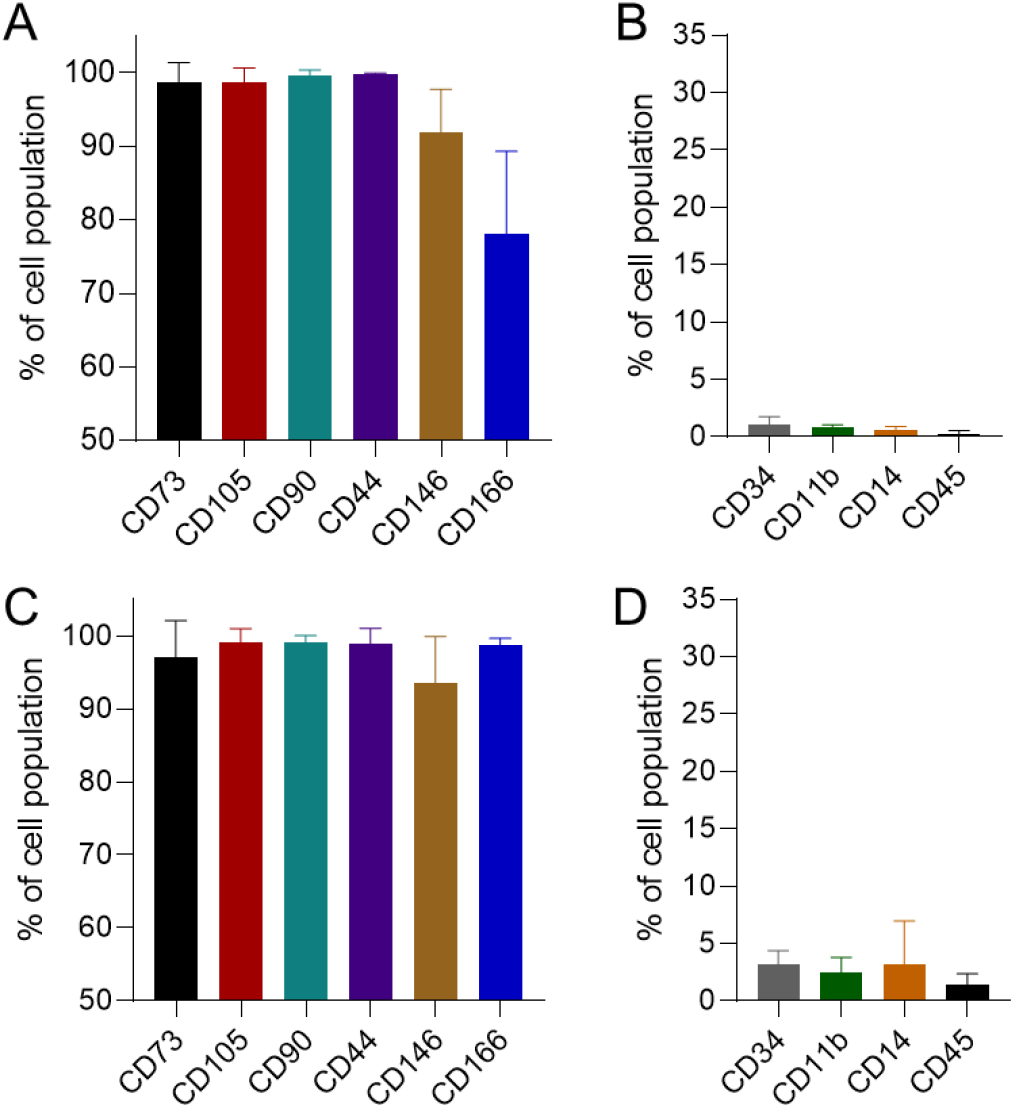
Expression profiles of cell surface antigens of BM-MSCs. Cells isolated and expanded to passages 2-3 either in xeno-/serum-free medium on the isoMATRIX (A, B) or in serum-containing medium on TCP (C, D) were examined by flow cytometry for the expression of cell surface markers CD73, CD105, CD90, CD44, CD146, CD166, CD11b, CD14, CD45 and CD34. The average percentage of cells positive for the indicated marker and the standard deviation is indicated. N = 6 (isoMATRIX), N = 5 (TCP)

### BM-MSCs isolated under xeno-/serum-free conditions using the isoMATRIX show significantly enhanced colony-forming efficiency, high trilineage differentiation potential and retain immunomodulatory capacity

A primary characteristic of MSCs is the ability to self-renew and form colonies. We evaluated the clonogenic potential of BM-MSCs (P0-1) using a colony-forming unit-fibroblast (CFU-F) assay. A total of 130 cells were seeded in a T75 flask, cultured for 14 days in the corresponding condition, and the number of colonies was counted after crystal violet staining. As shown in Fig. 3A-C, significantly more proliferative and adherent cells could be effectively isolated using the isoMATRIX combined with xeno-/serum-free medium (39 ± 7 colonies) compared to the serum-containing method (0.9 ± 1.5 colonies).

**Figure 3.**
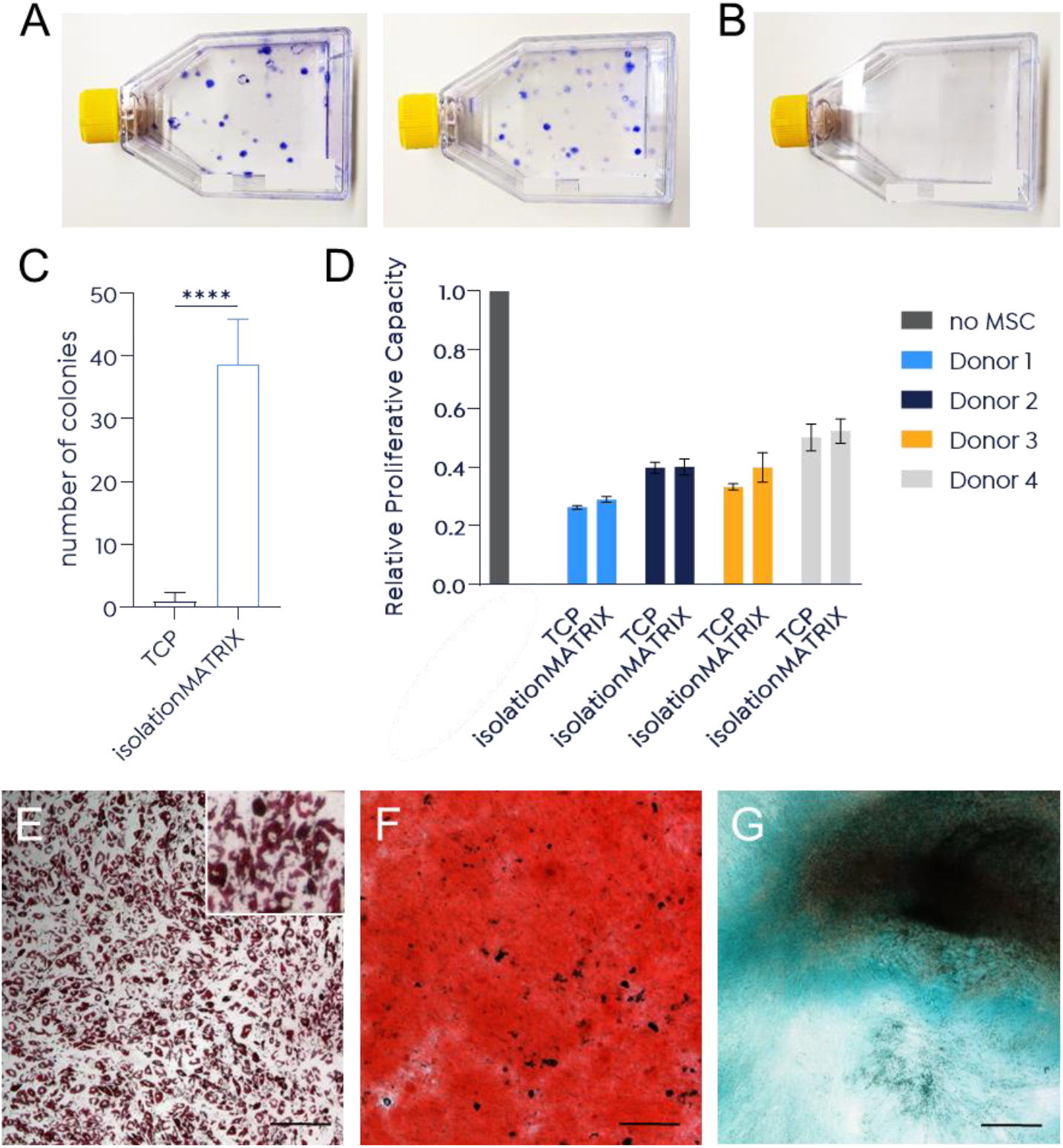
Enhanced colony-forming efficiency, high immunomodulatory capacity and trilineage differentiation potential of isoMATRIX-derived BM-MSCs. BM-MSCs isolated and expanded either in xeno-/serum-free medium using the isoMATRIX or in serum-containing medium on TCP were used to perform a CFU-F assay (A-C), a lymphocyte stimulation assay (D) as well as trilineage differentiation (E-G). A total of 130 cells (P0-1) were seeded in a T75 flask, cultured for 14 days in the appropriate condition (xeno-/serum-free medium on the isoMATRIX; duplicates are shown (A) or in serum-containing medium on TCP (B)) and stained with crystal violet for analysis (C). For the lymphocyte stimulation assay, cells (P2-3) were irradiated and incubated with CD3/CD28-stimulated PBMCs. The bars represent the relative proliferative capacity of cells displaying [3H]-thymidine incorporation compared to control (PBMC alone = 1.0) (D). For analysis of adipogenic, osteogenic and chondrogenic differentiation, cells were stained with Oil Red O for lipid droplets after 7 days (E), with Alizarin Red for calcium phosphate deposits produced by osteocytes after 28 days (F) and with Alcian Blue for proteoglycans synthesized by chondrocytes after 21 days of induction (G), respectively. Images were taken using Lionheart FX microscope (4x). Representative images are shown. N = 4. Scale bar, 200 μm.

To evaluate the immunomodulatory properties, we examined the impact of BM-MSCs on peripheral blood mononuclear cell (PBMC) proliferation. BM-MSCs of four different donors were used to perform a lymphocyte stimulation assay^26^. As demonstrated in Figure 3D, isoMATRIX-derived BM-MSCs were able to reduce the relative proliferative capacity to 0.3-0.5, similar to TCP-isolated cells. Thus, MSCs obtained from bone marrow aspirates under xeno-/serum-free isolation conditions maintain immunomodulatory capacities comparable to cells isolated under serum-containing conditions.

IsoMATRIX-derived BM-MSCs were also differentiated towards adipocytes, osteocytes, and chondrocytes to assess their in vitro trilineage differentiation potential. The adipogenic differentiation was visualized by staining cytoplasmic lipid droplets with Oil Red O after 7 days (Fig. 3E). Osteogenic and chondrogenic differentiation was observed by Alizarin Red staining of calcium phosphate deposits after 28 days and Alcian Blue staining of proteoglycans after 21 days, respectively (Fig 3F and 3G). All tested donors showed a high ability to differentiate into the three mesoderm-specific lineages.

In sum, our results demonstrate that the isolation of MSCs from bone marrow aspirates with the isoMATRIX under xeno-/serum-free conditions extracts a cell population with high proliferation and differentiation potential as well as strong immunomodulatory capacities.

### isoMATRIX improves the isolation efficiency of MSCs from adipose tissue in xeno-free and chemically defined medium

MSCs isolated from different tissue sources share a common phenotype (compare Fig. 1D and 4A), differentiation potential, and the expression of many cell surface markers, but differ in gene expression patterns, proliferation rate, and therapeutic efficacy^7–9^.

We performed an experiment using subcutaneous adipose tissue from a single donor and a different xeno-free medium and a chemically defined medium to exemplify the compatibility of the isoMATRIX with other tissue sources as well as medium compositions. Figure 4B shows that 6 days after isolation, the number of cells was markedly increased in both media on the isoMATRIX compared to the uncoated control (TCP) and serum-containing condition. The highest fold change (2.5-fold) in the number of isolated adipose tissue-derived MSCs (AD-MSCS) was observed using the chemically defined medium in combination with the isoMATRIX (CnT-PR-MSC-XF-HC + isoMATRIX: 8.15 ± 1.13 x 10^5^ cells versus TCP: 3.31 ± 0.3 x 10^5^). A significant increase in isolation efficiency was also seen under xeno-free conditions with a 1.7-fold increase in isolated cell number (CnT-PR-MSC-XF + isoMATRIX: 1.26 ± 0.06 x 106 cells versus TCP: 7.23 ± 0.08 x 10^5^). These results indicate that the isoMATRIX also enhances the isolation of MSCs from tissue sources other than bone marrow aspirates and underlines its compatibility with diverse media ranging from xeno-/serum-free to chemically defined.

**Figure 4.**
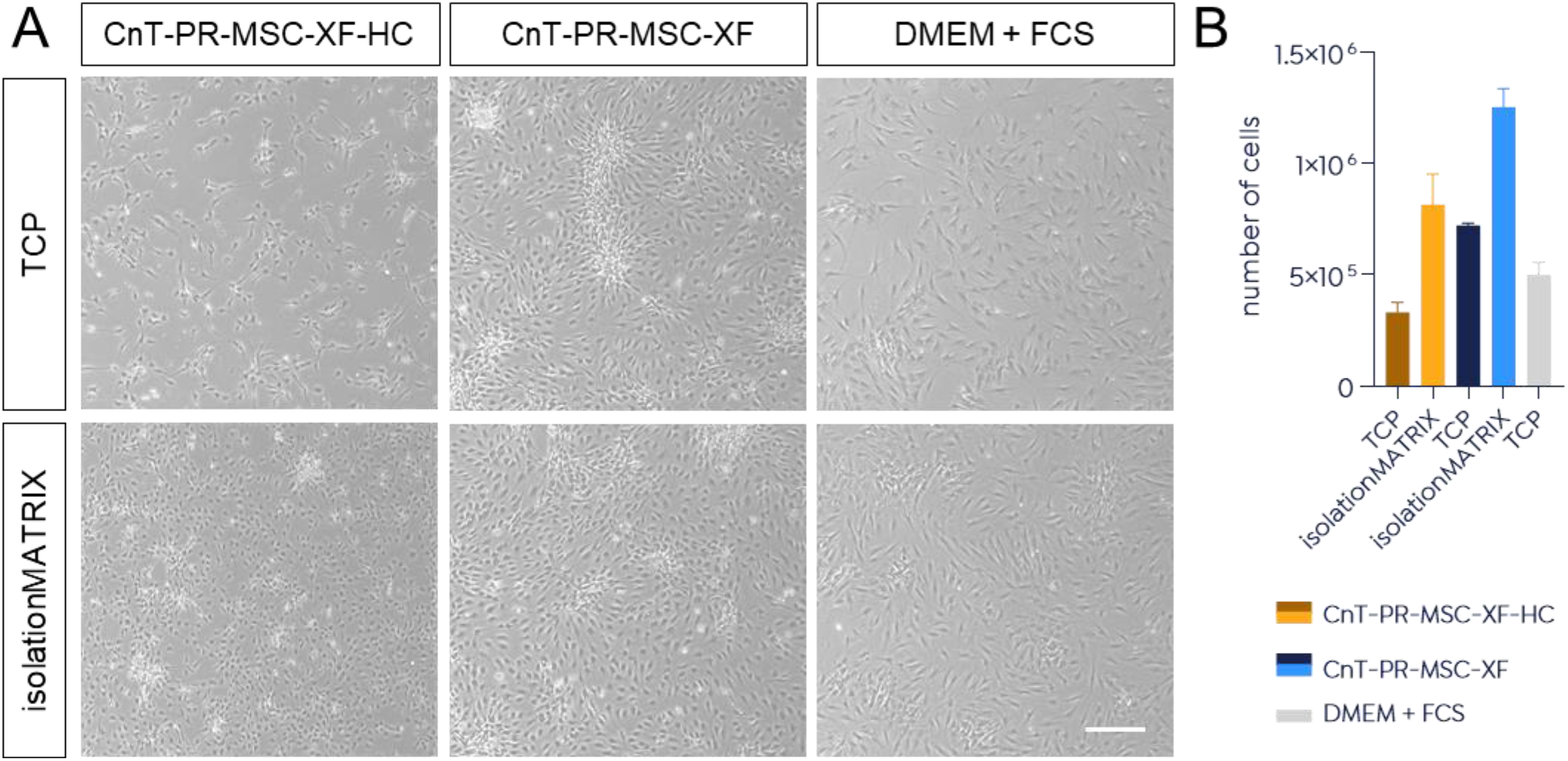
Isolation of MSCs from subcutaneous adipose tissue. Cells were isolated from the stromal vascular fraction of subcutaneous adipose tissue samples after enzymatic tissue digestion by seeding 20,000 cells/cm^2^ either on the isoMATRIX or TCP in combination with xeno-free (CnT-PR-MSC-XF), chemically defined (CnT-PR-MSC-XF-HC) or serum-containing medium (DMEM + FCS). Morphology (A) and cell number (B) were analyzed on day 6 post-isolation. N=1. Scale bar, 300 μm.

## Material and Methods

### Isolation and culture of MSCs from bone marrow aspirates

MSCs were isolated from healthy donors (n=10) after obtaining informed written consent (Ethical Approval Nos. EK221102004 and EK47022007). Briefly, human bone marrow aspirates were diluted in phosphate buffered saline (PBS, Thermo Fisher Scientific, Waltham, Massachusetts, United States) at a ratio of 1:5. A 25 mL aliquot was layered over a 1.073 g·mL^−1^ Ficoll-Paque™ PLUS solution (GE Healthcare, Chicago, Illinois, United States) and centrifuged at 2000 x g for 25 min at room temperature (RT). The mononuclear fraction was recovered, washed with PBS, and seeded as passage 0 either in Dulbecco’s Modified Eagle’s Medium (DMEM, low glucose, GlutaMAX supplement, pyruvate, Thermo Fisher Scientific) + 10% fetal calf serum (FCS; Sigma Aldrich, St. Louis, Missouri, United States) on plastic or in PRIME-XV MSC Expansion XSFM medium (FUJIFILM Irvine Scientific, Santa Ana, California, United States) on the isoMATRIX (denovoMATRIX GmbH, Dresden, Germany). Cells were cultured at 37°C at 5% CO_2_ in a humidified atmosphere and expanded further in the appropriate conditions. On day 4 after isolation, the cell number was determined by manually counting of 10×13 tile images (4X) using Fiji Software.^27^ Cells were harvested using TrypLE™ Express Enzyme (Gibco, Thermo Fisher Scientific) and subcultured in the same conditions. Viability, cell number and cell size were assessed using Trypan Blue dye exclusion with EVE cell counter (NanoEnTek, Seoul, South Korea). A full media change was performed twice a week.

### Flow cytometry

For the characterization of the MSC-specific surface phenotype, cells were incubated with fluorophore-coupled antibodies against CD11b, CD14, CD34, CD44, CD45, CD73, CD90, CD105, CD146 and CD166 or the corresponding isotype controls in PBS. For each sample, 30,000 – 50,000 events were acquired using Becton Dickinson (BD) fluorescence-activated cell sorting FACSCanto™ (BD Biosciences, Heidelberg, Germany). Histogram overlay subtraction analysis using FlowJo software (BD Biosciences) was performed to calculate the percentage of marker positive cells.

### In vitro multilineage differentiation assays

Cells were seeded in uncoated 24-well plates (serum-containing condition) or in those coated with myMATRIX MSC (xeno-/serum-free condition) at a density of 10,000 cells·cm^−2 20^. At 90% confluency, media were replaced by either complete adipogenesis (StemPro Adipogenesis Differentiation Basal Medium + Adipogenesis Supplement, Thermo Fisher Scientific) or osteogenesis medium (StemPro Osteocyte/Chondrocyte Differentiation Basal Medium + Osteogenesis Supplement, Thermo Fisher Scientific). Medium exchange was performed twice a week. For chondrogenic differentiation cell micromass cultures of 1.6 x 10^7^ cells were prepared and seeded as 5 μL droplets (=80,000 cells per droplet) in the center of each well. After 2 h of cultivation, complete chondrogenesis medium (StemPro Osteocyte/Chondrocyte Differentiation Basal Medium + Chondrogenesis Supplement, Thermo Fisher Scientific) was added and exchanged every 2-3 days. Adipogenic differentiation was stopped after 1-2 weeks, osteogenic differentiation after 4-5 weeks and chondrogenic differentiation after 3 weeks. All samples were rinsed once with PBS and fixed with 4% paraformaldehyde (PFA) for 10 min at RT. The extent of differentiation was determined microscopically, either by the appearance of Oil Red O-stained lipid vacuoles in adipocytes, Alizarin Red-stained calcium deposits produced by osteocytes, or Alcian Blue-stained proteoglycans synthesized by chondrocytes. Images were taken using Lionheart FX microscope (BioTek Instruments, Winooski, Vermont, United States, Gen5 software, version 3.03).

### CFU-F assay

130 cells (P0-1) were seeded in a T75 flask (TPP, Trasadingen, Switzerland) in the appropriate condition and cultured for 14 days. The corresponding medium (either DMEM+10% FCS or PRIME-XV MSC Expansion XSFM medium) was changed after 7 days of culture. After 14 days, the colonies were washed once with PBS and stained with 0.05 % crystal violet and counted.

### Lymphocyte Stimulation Assay

To test the immunomodulatory potential of isolated MSCs, a lymphocyte stimulation and subsequent thymidine incorporation assay was performed as described previously^26^. Briefly, PBMCs were isolated from healthy volunteer donors after obtaining written consent (Ethical approval No. EK206082008). The mononuclear cell fraction was isolated by Ficoll density gradient centrifugation, recovered, washed twice with PBS, and resuspended in Roswell Park Memorial Institute 1640 supplemented with 10% FCS. Lymphocyte stimulation was performed by mixing 1 x 10^5^ PBMCs, CD3/CD28 Dynabeads (Life Technologies, Carlsbad, United States) and 5 x 10^3^ irradiated (30 Gy, Gammacell 3000 Elan device, Best Theratronics Ldt., Ottawa, Canada) MSCs. After 5d of incubation at 37°C with 5% CO_2_ in a water-jacketed incubator, 1 μCi[3H]-thymidine (Hartmann Analytic, Braunschweig, Germany) was added to the culture. After an additional 18h incubation, cells were harvested and [3H]-thymidine incorporation was determined with the 1450 MicroBeta TriLux (PerkinElmer), converting degree of radioactivity to counts per minute (cpm).

### Isolation of adipose tissue-derived MSCs

Full thickness skin samples were obtained with informed consent of the donor according to the conditions listed in a valid ethics approval issued by the competent authority in the country of origin. Subcutaneous adipose tissue was minced and digested with collagenase I in PBS (2 mg/ml, Worthington, United States) at 37 °C for 90 minutes. The digest was diluted 1:1 with CnT-PR-MSC-XF-HC (CELLnTEC, Bern, Switzerland), a chemically defined, animal and human component-free culture medium, and the mix was let stand undisturbed to allow for phase separation. The lower aqueous phase was sequentially strained through 100 μm then 70 μm sieves. The resulting cell suspension was centrifuged, and the cell pellet was resuspended in 1 volume (V) CnT-PR-MSC-XF-HC before adding 3V of 1x Red Blood Cell Lysis Buffer (Biolegend, San Diego, United States). After 10 minutes of incubation at room temperature another 10V of CnT-PR-MSC-XF-HC were added and the cell suspension was centrifuged. The cell pellet was resuspended in CnT-PR-MSC-XF-HC and cells were counted. 500k cells per T25 cell culture flask were seeded in either of the following three culture media, CnT-PR-MSC-XF-HC, CnT-PR-MSC-XF (CELLnTEC) or DMEM (low glucose, GlutaMAX supplement, pyruvate) + 10% FCS in either uncoated or isoMATRIX-coated flasks. Medium was changed the next day, then every 2 to 3 days. On day 6 the cells were detached using CnT-Accutase-100 (CELLnTEC) and counted.

### Illustrations and statistical analysis

Illustrations were created with BioRender.com. Statistical analyses were performed using GraphPad Prism version 7.05 (GraphPad Software, La Jolla, United States). Outliers were evaluated using ROUT method (Q=1%). The significance of isolated cell numbers was assessed using the Mann-Whitney test. A two-tailed unpaired Student’s t-test was applied for the evaluation of average cell size and number of colonies in CFU-F assay. Data presentation and sample size (n) for each statistical analysis is indicated in the corresponding figure legend. Differences were regarded as significant if the calculated *p*-values were ≤0.05, *** = *p* < 0.005, **** = *p* < 0.0001.

## Discussion

MSC-based therapies are used to treat a variety of diseases, including Alzheimer’s disease^28–30^, bone and cartilage diseases^31–33^, autoimmune diseases^34,35^, diabetes^36,37^, graft-versus-host disease^38,39^, and multiple sclerosis^40,41^. MSCs offer many advantages for clinical applications such as easy accessibility, straightforward isolation and expansion procedures, high proliferative capacities, safety in autologous and allogeneic therapies, and the preservation of potency after storage. However, it is still challenging to manufacture an MSC-based therapeutic product with constant high yield and quality due to the lack of standardized procedures for harvesting, cultivation, and functional characterization as well as differences between donors and tissues of origin. Hence, setting up a reliable, consistent, and scalable system is critical in producing safe and potent MSCs on a broader scale despite the complex manufacturing process. Avoiding the use of serum-containing medium during both cell isolation and expansion is not only beneficial for product safety but helps to reduce lot-to-lot inconsistencies and enables commercialization of MSC-based cell therapies on a large scale. However, a xeno-/serum-free culture can reduce cell attachment and proliferation due to the lack of adhesion molecules, growth factors and/or nutrients in the medium. To support the use of xeno-/serum-free media without the need for human blood-derived supplements or animal-derived protein-coatings, we recently developed a chemically defined biomatrix (myMATRIX MSC) that promotes cell adhesion and supports the long-term expansion of MSCs with robust growth and high viability while maintaining their differentiation and immunomodulatory capacity, characteristic cell morphology, and the expression of key stemness markers^20^. To enable the isolation of MSCs under the same conditions, we developed a new matrix, called isoMATRIX, which is composed of dextran sulfate and an ECM protein-derived peptide-conjugate. Using the isoMATRIX in combination with the xeno-/serum-free PRIME-XV MSC Expansion XSFM medium (Fujifilm) for the isolation of MSCs from bone marrow aspirates resulted in an enhanced isolation efficiency of approximately 30% on day 4, which led to a 4-fold increase in the number of isolated cells on day 11-14 compared to serum-containing conditions. The cell population isolated with the isoMATRIX was more homogeneous and significantly smaller than serum-isolated controls but displayed the typical spindle-shaped, fibroblast-like morphology. The size of MSCs is often reported to correlate with their self-renewing properties and/or differentiation potential, with larger cells showing a lower proliferation and differentiation capacity and increased senescence^42–44^. Interestingly, Yin and colleagues showed that a continuous selection for a homogeneous and smaller population of MSCs results in faster proliferation and higher chondrogenic potential compared to conventional expansion procedures^45^. In our study, the smaller and more homogeneous cell population isolated with the isoMATRIX also showed a higher clonogenic and trilineage differentiation potential as well as a strong immunomodulatory capacity. In addition, isoMATRIX-derived MSCs expressed the characteristic profile of MSC-specific cell surface antigens. Like serum-isolated control cells, >95% expressed CD73, CD105, CD90, as postulated by the ISCT. They also showed no difference in the expression of CD44 and CD146. In contrast, we found elevated expression levels of CD34, CD11b, and CD14 in BM-MSCs isolated using TCP and serum-containing medium that even exceeded the limit of 2%, as defined by the ISCT. The expression of CD166 was reduced by ~20% in BM-MSCs isolated under xeno-/serum-free conditions using the isoMATRIX compared to the serum-containing condition. CD166 is a transmembrane protein of unknown function in MSC biology, and although significant variations have been observed for CD166 expression, it was identified as a possible human MSC gene expression or surface marker^46–49^. With the characterization assays that we have performed, we could not specifically link the reduction in CD166 surface expression to any of the MSC properties investigated. In sum, our results indicate that xeno-/serum-free isolation conditions extract a more uniform cell population and/or that the xeno-/serum-free microenvironment reduces cellular and metabolic heterogeneity that accumulates during in vitro expansion^50,51^.

Interestingly, isolation of MSCs from bone marrow aspirates with myMATRIX MSC, a surface coating optimized for MSC expansion, resulted in a suboptimal isolation efficiency (data not shown). Additionally, we were not able to isolate any cells using uncoated or fibronectin-coated culture vessels. The latter is recommended for MSC expansion using PRIME-XV MSC Expansion XSFM medium. After successful isolation, BM-MSCs can also be expanded in subsequent passages using the isoMATRIX with different media. However, their proliferation rates are significantly enhanced upon switching to the myMATRIX MSC instead (data not shown). These results illustrate the sensitivity of the cells towards their microenvironment and the strong impact of the provided artificial surface on different cell culture procedures (isolation or expansion). As discussed by Yuan and colleagues^10^, the transfer of MSC from their native microenvironment into non-physiological culture conditions with nutrient-rich medium and the lack of cell-cell-connections, cell-ECM-connections, different geometry, and mechanical properties results in a change of their energy metabolism from primarily glycolytic with active autophagy to a higher dependence on oxidative phosphorylation with reduced autophagy. This leads to increased senescence and reduced clinical potency of the cells as well as higher cellular and metabolic heterogeneity. Thus, a more accurate recreation of the natural microenvironment of human MSCs in vitro might lead to a more homogeneous and potent cell population. The superior isolation efficiency of the isoMATRIX highlights the benefit of a carefully tailored cell microenvironment.

Since MSCs can be isolated from various body tissues, we also tested the isoMATRIX using subcutaneous adipose tissue in combination with two different media. Irrespective of the medium, the isoMATRIX markedly increased cell yield in both conditions compared to uncoated controls with a fold increase in isolated cell numbers ranging from 1.7 to 2.5. These results indicate that the isoMATRIX can significantly enhance the isolation efficiency of MSCs using diverse media and different tissue sources.

Depending on the site of origin, there is a substantial difference in the number of MSCs, nevertheless, they are a rare population of progenitors in adult tissues. Consequently, the quantity of cells obtained after isolation is often insufficient for a clinical dose of about 10^6^-10^9^ cells^52,53^. This problem is potentiated by a decrease in growth kinetics, differentiation ability, and potency of MSC during extensive expansion in vitro. Hence, efficient MSCs isolation is the premise for optimized expansion procedures required to produce high-quality cells in clinically relevant numbers. The isoMATRIX in combination with a xeno-/serum-free medium provided on average a 30% increase in cell number after isolation. To investigate the impact of enhanced isolated cell numbers on the manufacturing process of an MSC-based therapeutic product, we have modeled an expansion process for manufacturing a therapeutical dose of 10^7^ cells/kg of body weight for a patient of 80 kg (Fig. 5A, reference procedure). Our model is based on a growth curve of MSCs expanded on the isoMATRIX in xeno-/serum-free medium (Fig. 5B). Starting the expansion process with 30% more isolated cells would increase cell production efficiency from 1-2 doses to 4 doses in the same time frame. This increase is amplified by the fact that the cell growth is roughly 3-7 times faster on the isoMATRIX in the xeno-/serum-free medium than in serum-containing conditions (Fig. 5B-C). This improvement has several implications for the manufacturing process of human MSCs for autologous and allogeneic therapies (Fig. 5A). Autologous therapies are often restricted by the patient's low MSC frequency and/or poor quality due to age or a diseased state. Hence, increasing the isolation efficiency accompanied by constant or even enhanced cell proliferation enables the treatment of the patient in a shorter time. In allogeneic stem cell therapy, cells from one donor are utilized for multiple patients and provided as an off-the-shelf product. In such therapies, the higher number of isolated cells obtained with the isoMATRIX at the beginning of a manufacturing process can substantially increase the final cell yield that is naturally limited by the lifetime of MSCs. In our model, an increase of 30% in starting material would result in sufficient cells to treat not only 1-2 patients but four patients. This impact would be potentiated in large-scale manufacturing, which will help reduce costs and expand the access of MSC-based cell therapy to many patients. Thus, our chemically defined biomatrix combined with xeno-, serum-free, or chemically defined media, promotes enhanced isolation of human MSCs and supports consistent and reliable cell performance for improved stem cell-based therapies.

**Figure 5.**
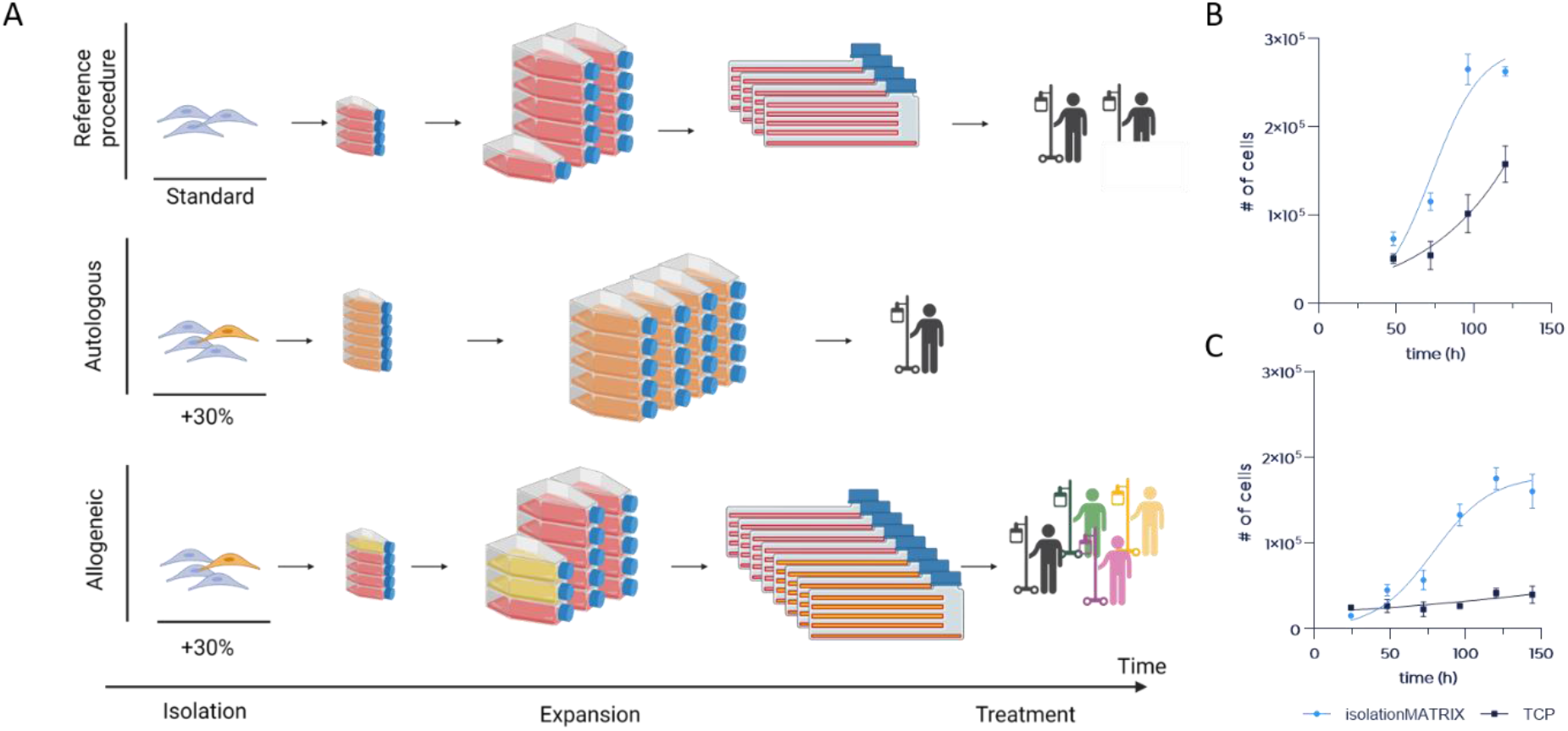
Modeling the expansion process for an MSC-based therapeutical product. The expansion process for a therapeutical dose of 10^7^ cells/kg for a patient of 80 kg was modeled assuming a seeding density of 5,000 cells/cm^2^ and is illustrated in A. The manufacturing starts with an isolated cell population that is propagated by stepwise surface enlargement until administration of cells into a patient. For autologous therapies, a 30% increase in starting material in combination with a slightly reduced seeding density (3,000 cells/cm^2^) would shorten manufacturing time. The impact of an 30% increase in starting material on the manufacturing process of human MSCs for allogeneic therapies while maintaining the seeding density of 5,000 cells/cm^2^ is indicated in yellow/orange. The expansion process model is inferred from a growth curve of MSCs expanded on the isoMATRIX in xeno-/serum-free medium (B). A second, slower proliferating donor is shown for comparison (C). Both donors were also cultivated on TCP in serum-containing medium over a time period of 5-6 days for comparison with standard expansion procedures. The growth curve data was fitted to the logistic growth model (Y=YM*Y0/((YM-Y0)*exp(-k*x) +Y0; nonlinear fitting) with R^2^= 0.91 (isoMATRIX, A), R^2^= 0.87 (TCP, A), R^2^= 0.96 (isoMATRIX, B) and R^2^= 0.71 (TCP, B).

## Acknowledgment

denovoMATRIX GmbH was supported and received funding by the European Social Fund (ESF), the European Regional Development Fund (ERDF), and by the EXIST-Forschungstransfer granted by the BMWi.

